# Longitudinal analysis of Flower-dependent cell competition fitness markers in *Drosophila melanogaster*

**DOI:** 10.1101/2024.03.25.586630

**Authors:** Mariana Marques-Reis, Barbara Hauert, Eduardo Moreno

**Affiliations:** Cell Fitness Lab, Champalimaud Centre for the Unknown, Av. Brasília, 1400-038 Lisbon, Portugal; Institute for Cell Biology, University of Bern, Baltzerstrasse 4, 3012 Bern, Switzerland

**Keywords:** Cell Competition, Flower LoseB, Azot, Apoptosis

## Abstract

Cell competition is a conserved phenomenon spanning from arthropods to humans. It involves the elimination of viable yet suboptimal “loser” cells when juxtaposed with their fitter “winner” counterparts. This process has received increased attention for its implications in cancer initiation and progression, neurodegeneration, and ageing.

This study investigates the presence of the loser fitness fingerprint Flower LoseB (Fwe LB) and the fitness checkpoint Azot in the optic lobes over a period of 28 days. Notably, the absence of Azot is conventionally linked to the accumulation of loser cells over time. However, our investigation reveals that this accumulation is not perpetual and, intriguingly, Azot is not required for loser cell elimination in this context. Furthermore, we estimate that fewer than 50% of Fwe LB-expressing cells also express Azot and undergo apoptosis. Remarkably, our calculations also demonstrate that over 50% of cells undergoing apoptosis at any given time point are positive for the loser markers Fwe LB and Azot.

This comprehensive analysis of fitness marker dynamics over a 28-day timeframe sheds new light on the intricate mechanisms governing Flower-dependent cell competition.

## Introduction

The fundamental principle of “survival of the fittest,” a cornerstone of Darwinian evolution, extends its relevance from the macroworld to the cellular level through a phenomenon known as cell competition. First observed in 1975, this phenomenon was characterised by the elimination of slower proliferating clones, specifically those carrying the ribosomal *minute* mutation, when near their wild-type counterparts (Morata and Ripoll 1975). Subsequent studies refined the concept, highlighting that not all disparities in cell proliferation rates trigger competition (Böhni et al. 1999; De La Cova et al. 2004), ultimately defining cell competition as the selective elimination of viable yet suboptimal cells in the presence of fitter counterparts within the same tissue compartment (Moreno and Rhiner 2014).

This broadly defined process has been documented in various organisms, including *Drosophila melanogaster* (Moreno, Basler, and Morata 2002), zebrafish (Walderich et al. 2016), mice (Oliver et al. 2004), and humans (Madan et al. 2019). It has been implicated in various biological processes, from cancer initiation and proliferation to brain injury, senescence, stem cell niche dynamics, neurodegeneration, and ageing (Bondar and Medzhitov 2010; Coelho et al. 2018; Coelho and Moreno 2020; Fernández-Hernández, Rhiner, and Moreno 2013; Marques-Reis and Moreno 2021; Merino et al. 2015; Moreno 2008; Moreno et al. 2015; Rhiner et al. 2009). Cell competition can be categorised into three principal causative factors: competition for survival factors, where cells in closer proximity to these factors gain a competitive advantage (Moreno et al. 2002); competition for spatial occupation, wherein mechanically less resilient cells are outcompeted (Levayer, Dupont, and Moreno 2016; Moreno et al. 2019); and competition based on fitness differences, leading to the elimination of suboptimal cells, potentially replaced by fitter counterparts (Merino, Levayer, and Moreno 2016). Numerous proteins have been implicated in regulating the winner or loser status of a cell, including Flower (Rhiner et al. 2010), Slit-Robo-Ena (Vaughen and Igaki 2016), Spätzle-Toll (Alpar, Bergantiños, and Johnston 2018; Germani et al. 2018; Meyer et al. 2014), Sas-PTP10D (Yamamoto et al. 2017), and Flamingo (Bosch, Cho, and Axelrod n.d.). The focus of this work centres exclusively on Flower-dependent cell competition.

Flower, a transmembrane protein with predicted calcium channel activity, plays a central role in cell competition (Rhiner et al. 2010; Yao et al. 2009). Nevertheless, its predicted calcium channel function does not appear to influence the competitive process (Coelho and Moreno 2020; Madan et al. 2019). In *Drosophila*, Flower has three distinct isoforms -Flower Ubi, predominantly present in winner cells, and Flower LoseA and Flower LoseB (Fwe LB), traditionally associated with loser cells (Rhiner et al. 2010). Flower is conserved in mice and humans, where it has been implicated in contexts such as cancer development and poor prognosis in COVID-19 cases (Madan et al. 2019; Petrova et al. 2012; Yekelchyk et al. 2021).

After being labelled as such, loser cells must pass additional checkpoints before their elimination, as not all loser cells are eliminated (Rhiner et al. 2010). In *Drosophila*, loser cells not meant for elimination express *SPARC*, which prevents their elimination (Portela et al. 2010). Conversely, loser cells marked for elimination express *azot*, producing a four-EF-hand cytoplasmic protein responsible for inducing the pro-apoptotic gene *hid* (Merino et al. 2015). Azot is a recognised prerequisite for Flower-dependent cell elimination, as its absence results in the accumulation of loser cells with time (Merino et al. 2015). This depletion of *azot* correlates with a diminished lifespan and an increased prevalence of degenerative vacuoles (Merino et al. 2015). Notably, alterations in *azot* expression in the gut correlate with corresponding changes in lifespan, reinforcing the pivotal role of Azot and cell competition in the ageing process (Merino,2023).

The primary objective of this study was to study the presence of Fwe LB and Azot over a 28-day timeframe in the optic lobes, within contexts where Flower-dependent cell competition occurs normally contrasted with situations where the absence of *azot* compromised competition. Our findings unveil that, in the absence of *azot*, loser cells do not accumulate in perpetuity.

Intriguingly, even in the absence of *azot*, loser cells continue to be eliminated, indicating that Azot is not an absolute requisite for loser cell elimination in this context. Additionally, our investigation sought to elucidate the proportions of loser cells that undergo apoptosis and, reciprocally, the percentages of dying identified as loser cells, as not all loser cells are eliminated, and not all dying cells are losers (Merino et al. 2015; Rhiner et al. 2010).

This work contributes to the comprehensive understanding of fitness marker expression throughout time while shedding light on the contribution of cell competition in shaping the landscape of cell elimination. Moreover, it introduces the concept of compensatory mechanisms within animals to mitigate loser cell persistence when the canonical Flower-dependent cell competition process is compromised.

## Results

### Accumulation of loser cells in an *azot* knockout scenario is not perpetual

In this study, we generated a transgenic *Drosophila melanogaster* to concomitantly mark the presence of Fwe LB and Azot. Specifically, Azot labelling was accomplished by replacing one copy of the *azot* gene with a LexA::p65 fusion protein (Figure *1*A), which activates a LexAOP driving the expression of CD8::GFP under the *azot* promoter. The use of this genetic tool was based on prior research indicating that a single copy of the *azot* gene is sufficient for its functional role (Merino et al. 2015).

**Figure 1.**
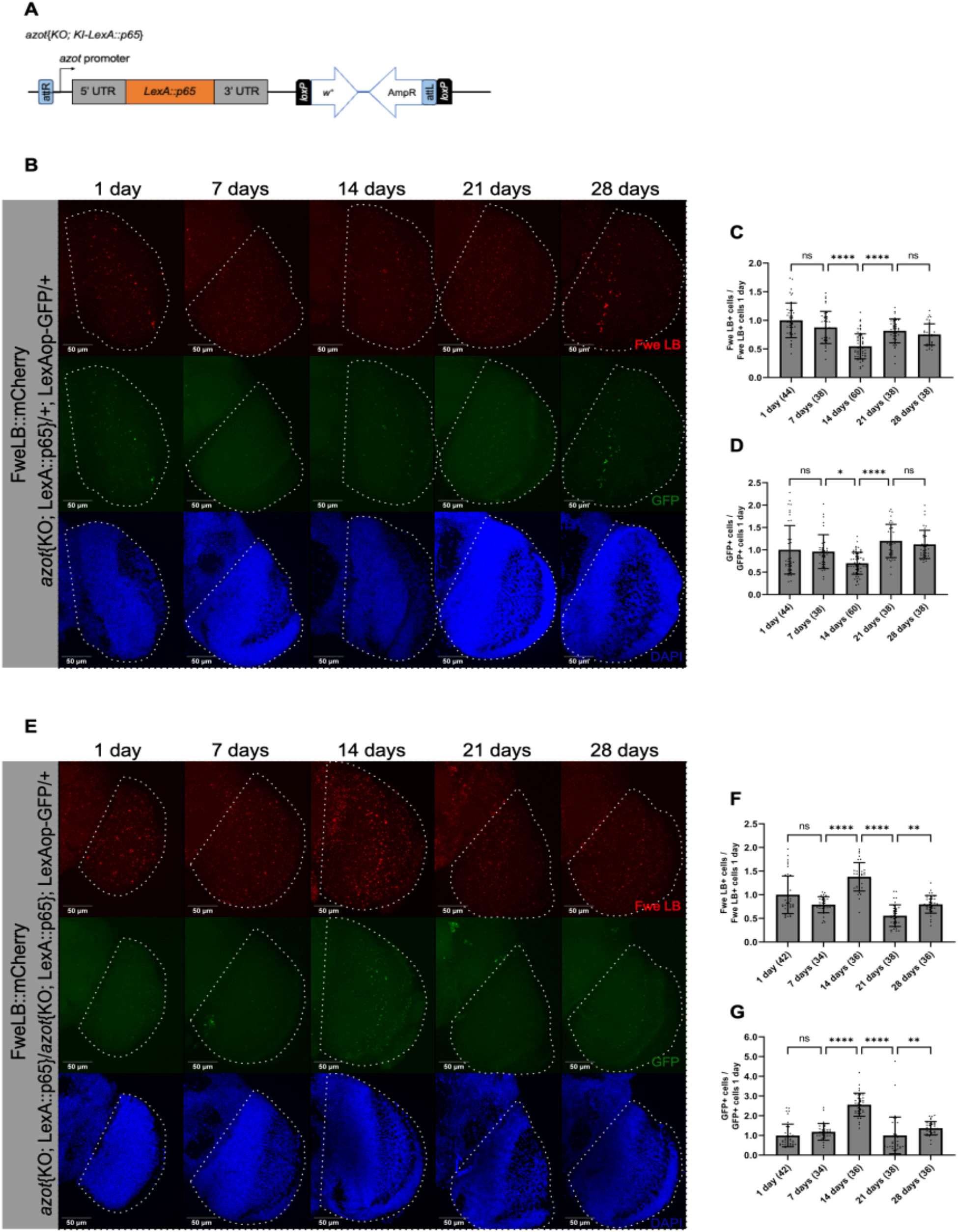
Expression of the fitness markers Flower LoseB and Azot over 28 days. (A) Scheme of the modified *azot{KO;KI-LexA::p65}* locus. This transgenic line was generated by integration of a knockin construct containing the LexA::p65 sequence under the control of the endogenous *azot* promoter, into the *azot* knockout locus. The vector backbone (*w*^*+*^, *AmpR*) was maintained in the knockin line. (B) Adult optic lobes of flies with one copy of *azot*, 1, 7, 14, 21 and 28 days old. Fwe LB is represented in red, GFP in green and DAPI in blue. Scale bars, 50 μm. Quantification of Fwe LB positive cells (C) or GFP (D) in the optic lobe of flies with one copy of *azot*, 1, 7, 14, 21 and 28 days old normalized against their respective numbers at 1 day old. (E) Adult optic lobes of flies azot KO, 1, 7, 14, 21 and 28 days old. Fwe LB is represented in red, GFP in green and DAPI in bluei. Scale bars, 50 μm. Quantification of Fwe LB positive cells (F) or GFP (G) in the optic lobe of flies azot KO, 1, 7, 14, 21 and 28 days old normalized against their respective numbers at 1 day old. The numbers after the age of the flies indicate the number of optic lobes analyzed. Error bars indicate SD; NS indicates non-significant; *P<0.05; **P<0.01; ****P<0.0001. Statistical significance between groups was calculated using the nonparametric Kruskal-Wallis test and a Dunn’s test was applied for multiple comparisons between genotypes. Genotypes: ywF; *azot*{KO; KI-*LexA::p65*}/+; 26xLexAop-CD8::GFP, *flower*{KO; KI-*flowerLoseB::mCherry*}/+ (A-C). ywF; *azot*{KO; KI-*LexA::p65*}/ azot{KO; KI-*LexA::p65*}; 26xLexAop-CD8::GFP, *flower*{KO; KI-*flowerLoseB::mCherry*}/+ (D-F).

Employing this genetic tool, we were able to monitor the temporal manifestation of both Fwe LB and Azot over a period of 28 days, with observations conducted at seven-day intervals (Figure *1*B). Intriguingly, our findings revealed a noteworthy reduction in the number of Fwe LB-positive and GFP-positive cells at the 14-day time point (Figure *1*C-D).

Furthermore, this tool enabled us to assess the numbers of Fwe LB and Azot-positive cells in a genetic context where both copies of the *azot* gene were substituted with LexA::p65 – *azot* knockout (KO) -(Figure *1*E). Our analysis revealed a 1.4-fold increase in the number of cells positive for Fwe LB and a 2.6-fold increase in the number of cells positive for GFP at the 14-day time point, compared to their respective levels at one day old. This observation aligns with prior findings by Merino et al. 2015, suggesting the accumulation of cells attempting to express Azot, and additionally reveals a parallel increase in the population of Fwe LB-expressing cells (Figure *1*F-G).

Surprisingly, at the subsequent time point of 21 days, there was a decline in Fwe LB-positive cells to 0.6-fold and in GFP-positive cells to 1.0-fold, relative to their respective levels at one day old. These data contravene the view that accumulating “loser” cells in an *azot* KO scenario is perpetual and led us to hypothesise that these cells may undergo apoptosis.

### Azot is not required for loser cell elimination in a time-dependent scenario

To investigate whether loser cells in an *azot* KO scenario die, we used an antibody against Dcp1, a marker indicative of caspase activation. This analysis revealed the presence of apoptotic cells among the population lacking *azot* (as illustrated in Figure *2*A). Notably, our data unveiled a peak in the population of cells simultaneously positive for Fwe LB, Azot, and Dcp1 at the 14-day stage (Figure *2*B), strongly suggesting that the reduction in the number of loser cells within the optic lobes at 21 days is closely associated with the peak of apoptotic elimination of these cells at the 14-day time point.

**Figure 2.**
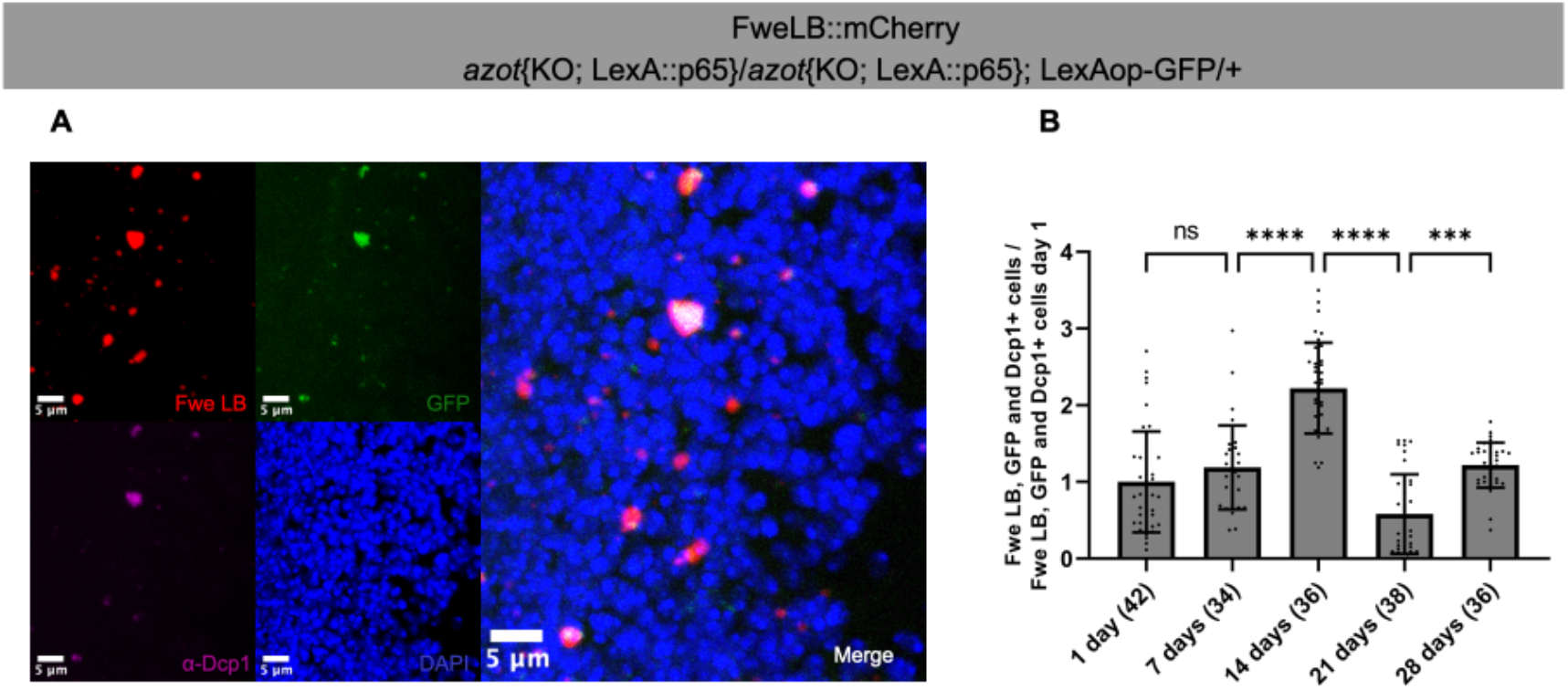
Loser cells die in the absence of *azot*. (A) Detail of an optic lobe of flies *azot* KO at 14 days old. Fwe LB is represented in red, GFP in green, α-Dcp1 in magenta, and DAPI in blue. Scale bars, 5 μm. (B) Quantification of cells co-labelled with FweLB, GFP and α-Dcp1 at 1, 7, 14, 21 and 28 days old normalized against their number at 1 day old, in optic lobes *azot* KO. The numbers after the age of the flies indicate the number of optic lobes analyzed. Error bars indicate SD; NS indicates non-significant; ***P<0.001; ****P<0.0001. Statistical significance between groups was calculated using the nonparametric Kruskal-Wallis test and a Dunn’s test was applied for multiple comparisons between genotypes. Genotype (A-B): ywF; *azot*{KO; KI-*LexA::p65*}/ *azot*{KO; KI-*LexA::p65*}; 26xLexAop-CD8::GFP, *flower*{KO; KI-*flowerLoseB::mCherry*}/+.

These findings demonstrate that loser cells without *azot* are indeed subject to elimination through apoptosis. Consequently, our data firmly establish that the depletion of *azot* alone is insufficient to impede the Flower-dependent process of cell elimination, at least in a time-dependent scenario.

### Being labelled as loser is not a death sentence, and not all dying cells are loser

In the context of homeostatic Flower-dependent cell competition, which occurs in animals possessing functional Fwe LB and Azot proteins, and considering the hierarchical relationship between them, it is anticipated that only a subset of Fwe LB-positive cells will simultaneously express Azot. Furthermore, only a fraction of the Azot-positive cells are expected to undergo cell death. By marking the cells co-expressing Fwe LB, Azot, and the Dcp1, we were able to quantify the proportion of loser cells (those expressing both Fwe LB and Azot) that undergo apoptosis in relation to the overall population marked with the loser fitness fingerprint Fwe LB or in relation to the overall population marked with Dcp1.

Our analysis revealed that the proportion of cells marked with Fwe LB, expressing Fwe LB, Azot, and Dcp1, remained relatively constant over time, with average values fluctuating between 33.8% and 50.0% (Figure *3*A). This observation implies that less than half of the cells labelled with the loser fitness fingerprint Fwe LB actively engage in the Flower-dependent cell competition pathway by expressing Azot and ultimately dying.

**Figure 3.**
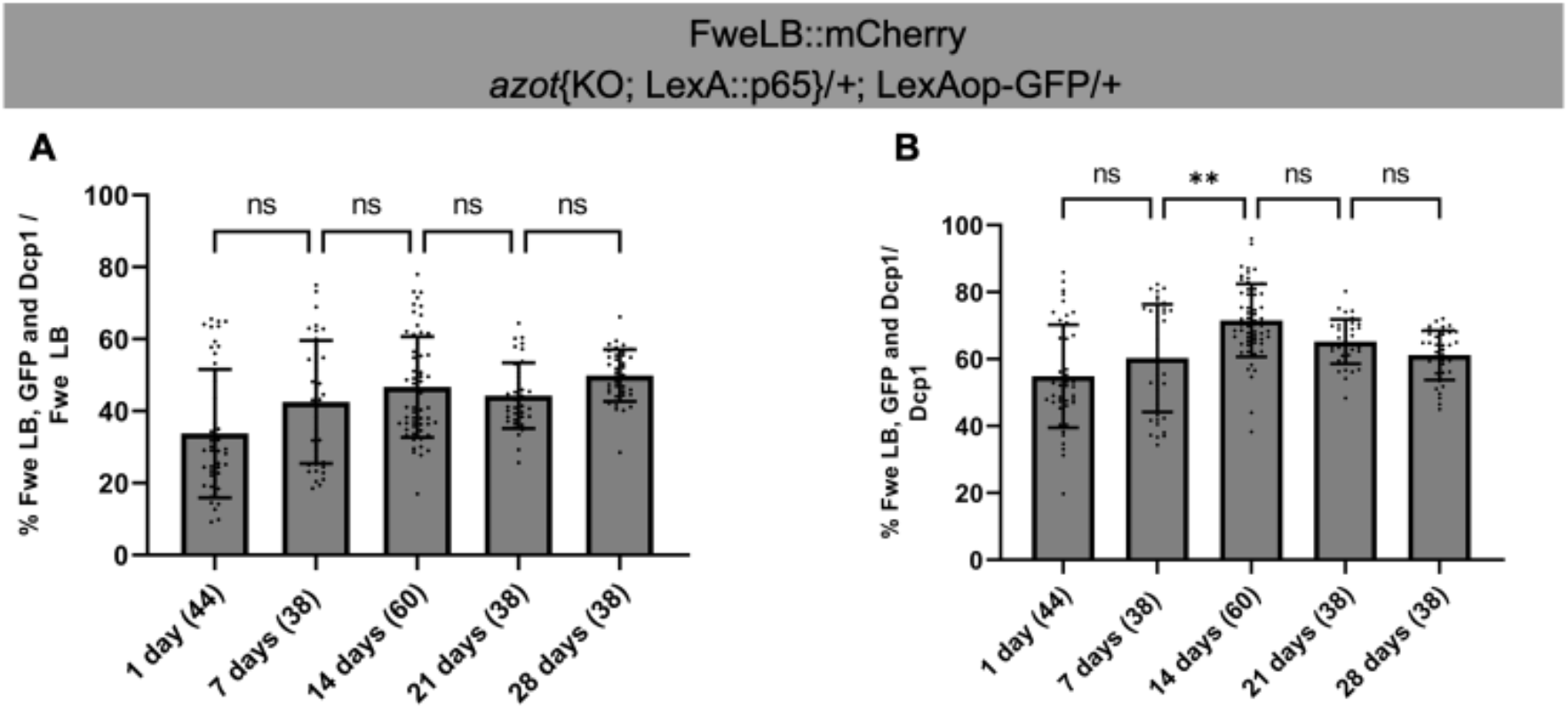
Percentage of loser cells dying over the total number of loser or dying cells. (A) Percentage of cells co-labelled with FweLB, GFP and α-Dcp1 over the total number of Fwe LB positive cells, at 1, 7, 14, 21 and 28 days old, in optic lobes of flies with one copy of *azot*. (B)Percentage of cells co-labelled with FweLB, GFP and α-Dcp1 over the total number of Dcp1 positive cells, at 1, 7, 14, 21 and 28 days old, in optic lobes of flies with one copy of *azot*. The numbers after the age of the flies indicate the number of optic lobes analyzed. Error bars indicate SD; NS indicates non-significant; **P<0.01. Statistical significance between groups was calculated using the nonparametric Kruskal-Wallis test and a Dunn’s test was applied for multiple comparisons between genotypes. Genotype (A-B): ywF; *azot*{KO; KI-*LexA::p65*}/ +; 26xLexAop-CD8::GFP, *flower*{KO; KI-*flowerLoseB::mCherry*}/+.

We also calculated the percentage of cells positive for Fwe LB, Azot, and Dcp1 in relation to the total number of cells undergoing apoptosis. Over the course of the experiment, this average percentage varied between 55.0% to 71.6% (Figure *3*B). This outcome indicates that more than half of the cells undergoing apoptosis in the optic lobe are expressing “loser” fitness markers, emphasising the significance of the Flower-dependent cell competition process in shaping cellular fate in this region.

## Discussion

Over time, brain cells can become suboptimal due to wear and tear, excitotoxicity, reactive oxygen species-induced damage, or hypoglycemic episodes (Hutchins and Barger 1998). It is conceivable that the decrement in the functional state of certain cells precipitates their categorisation as loser. Consequently, the upsurge in the quantity of Fwe LB and Azot-positive cells from day 14 to day 21 may be attributed to the natural ageing process. However, it is intriguing to note that at 1 and 7 days, the count of Fwe LB and Azot-positive cells is higher compared to 14 days. Observations from murine models suggest that approximately 50% of neurons generated are eliminated during the first postnatal week (Castillo-Ruiz et al. 2020). In the case of *Drosophila*, neurons participating in the ecdysis process experience rapid elimination within 24 hours following eclosion (Kimura and Truman 1990; Pinto-Teixeira, Konstantinides, and Desplan 2016; Tissot and Stocker 2000). Analogous to the elimination of supernumerary ommatidia during the pupal stage, which is known to be Flower-dependent (Merino et al. 2013), it is plausible that certain cells within the optic lobes, without function in the adult state, may also undergo elimination via the process of cell competition, explaining the high numbers of Fwe LB and Azot-positive cells at 1 and 7 days. Subsequent research is warranted to substantiate this idea.

In the *azot* KO optic lobes, the initial accumulation of Fwe LB or GFP-positive cells, indicative of those that would express Azot, within the first 14 days, followed by subsequent diminishment, unveils a compelling temporal dynamic that warrants further investigation. Our data shows that those loser cells, marked by Fwe LB and GFP, undergo apoptosis, with a noticeable peak observed at 14 days. We propose that during the initial 14-day period, the pace of loser cell labelling surpasses their elimination rate, culminating in the accumulation of loser cells. At the 14-day mark, this equilibrium between accumulation and elimination is disrupted, leading to a surge in loser cell apoptosis. Intriguingly, the timeframe spanning from day 21 to day 28 exhibits a resurgence in the accumulation of loser cells, concomitant with a reduction in their apoptosis compared to the 14-day mark. This observation suggests the cyclic nature of loser cell accumulation and elimination in the absence of *azot*. It is plausible that, in the absence of Azot, this process of accumulation and elimination may not be optimised, which aligns with the understanding that flies lacking Azot have reduced lifespans (Merino 2023; Merino et al. 2013). This observation may indicate the presence of compensatory mechanisms employed by flies to eliminate suboptimal cells when the canonical Flower-dependent cell competition process is compromised, akin to the redundancy of caspases such as Dcp1 and drlCE in certain cellular contexts (Steller 2008; Xu et al. 2009), which ensure the seamless execution of apoptosis.

Notably, fewer than 50% of Fwe LB-positive cells express Azot and subsequently undergo elimination, confirming that not all loser cells proceed along this trajectory towards elimination. From their labelling as a loser, cells can express SPARC (Portela et al. 2010), which protects them from elimination. Moreover, if loser cells do not have fitter neighbours, elimination does not occur (Rhiner et al. 2010). Another intriguing observation is that over 50% of cells undergoing apoptosis exhibit the loser markers, Fwe LB and Azot. This underscores the active role of Flower-dependent cell competition in orchestrating most of the cell death in the optic lobes, aligned with Coelho et al. 2018 findings.

It is imperative to acknowledge a limitation within our experimental design. The cross-sectional nature of our study, characterised by data collection at discrete time points, may not entirely encapsulate the continuous dynamics governing the labelling and elimination of loser cells. This design precludes the possibility that certain Fwe LB-positive cells, which had not yet expressed Azot at the time of observation, may subsequently undergo this transition. This circumstance suggests that the calculated percentages may underestimate the overall prevalence of this process and reinforces the need for further investigations.

## Conclusion

The findings discussed in this study shed light on the complex dynamics of cell competition within the *Drosophila* optic lobes over time. In the absence of *azot*, the temporal dynamics observed, from the initial accumulation of loser cells to their subsequent elimination, followed by a resurgence in accumulation, challenge prior assumptions about the linear nature of this process and suggest the fly has redundant mechanisms to eliminate loser cells when the canonical Flower-dependent cell elimination process is compromised. The cyclical nature of loser cell dynamics and the interplay between the pace of loser cell labelling and their elimination in the absence of *azot* emerge as compelling areas for future investigation. Furthermore, the active role of Flower-dependent cell competition in orchestrating cell death shows its relevance in the cell homeostasis of the optic lobes. The intricacies of cell competition in the *Drosophila* optic lobes offer a rich field for further exploration and promise to deepen our understanding of the mechanisms governing cell fitness and survival in complex biological systems.

## Experimental Procedures

### Drosophila Genetics

Stocks and crosses were kept at 25ºC in Vienna standard media with extra yeast. Flies were collected after eclosion and dissected at 1, 7, 14, 21, and 28 days old. The following stocks were used: ywF; *azot*{KO; KI-*LexA::p65*}/CyO; 26xLexAop-*CD8::GFP, flower*{KO; KI-*flowerLoseB::mCherry*}/TM6B and ywF; *azot*{KO; KI-*LexA::p65*}/CyO;.

### Azot knockin Generation

The *azot* knockout founder line was described by Merino et al. 2015. For the generation of the *azot*{KO; KI-*LexA::p65*}, the cDNA of LexA::p65 was generated and inserted into a vector *w*^*+*^, *AmpR*, and the knockin (KI) was generated as described in (Huang et al. 2009). Primer sequences are available upon request.

### Immunohistochemistry and image acquisition

Adult brains were dissected in cold PBS; the samples were fixated for 30min in formaldehyde (4% v/v in PBS) and permeabilised with PBT 1% Triton. The brains were then incubated with rabbit α-Dcp1 (1:50) from Cell Signaling (#9578). Samples were mounted in Vectashield with DAPI (Vectorlabs), and the confocal images were acquired with Zeiss LSM 880 using the Plan-Apochromat 40x/1.4 Oil DIC M27 objective. Maximum intensity projections of the 71-μm-wide images were obtained with Zeiss Zen Black.

### Quantification and statistical analysis

The number of Fwe LB, GFP, and Dcp1 positive cells and colocalisation were quantified with a Fiji macro developed for this work, available upon request. A minimum number of 30 optic lobes were analysed for each condition. Statistical analysis was performed using GraphPad Prism 9.

Statistical significance between groups was calculated using the nonparametric Kruskal-Wallis test, and Dunn’s test was applied for multiple comparisons between genotypes.

## Abbreviations

Fwe LB: Flower LoseB
KI: knockin
KO: knockout

## Acknowledgements

We thank Bloomington Stock Center for flies; the technicians at the Champalimaud Fly Platform for support with stock maintenance; the ABBE platform for microscopy support; Catarina Brás-Pereira for her critical feedback; Andrea Spinazzola and Andrés Gutiérrez for their suggestions and comments on the manuscript. M.R. was supported by an FCT -Fundação para a Ciência e a Tecnologia - PhD studentship (SFRH/BD/138537/2018). Portuguese national funds supported this study through FCT in the context of the project UIDB/04443/2020 and the European Research Council (Consolidator Grant to E.M.: ‘‘Active Mechanisms of Cell Selection: From Cell Competition to Cell Fitness’’). Fly platform was funded by the research infrastructure CONGENTO, co-financed by Lisboa Regional Operational Programme (Lisboa2020), under the PORTUGAL 2020 Partnership Agreement, through the European Regional Development Fund (ERDF) and Fundação para a Ciência e Tecnologia (Portugal) under the project LISBOA-01-0145-FEDER-022170. The Portuguese Platform of Bioimaging funded ABBE platform - LISBOA-01-0145-FEDER-022122.

## Notes

### Competing Interest Statement

The authors have declared no competing interest.

